# A systematic approach for finding herbicide synergies

**DOI:** 10.1101/2021.02.08.430187

**Authors:** Kirill V. Sukhoverkov, Joshua S. Mylne

## Abstract

Combining herbicides into a double dose is a common approach to overcome the potential for herbicide resistance by weeds. Many herbicide mixtures can be antagonistic and they are rarely synergistic. Here, 24 commercial herbicides, each representing a mode of action were used to create a matrix of all 276 unique combinations to search for new synergies in agar with *Arabidopsis thaliana*. Herbicides were used at an appropriate sub-lethal dose such that any synergies gave visible growth inhibition. We found five synergies including three new ones, namely mesotrione-norflurazon, mesotrione-clethodim and paraquat-clomazone. All three new synergies were reproducible in soil-grown conditions. Interestingly, all three new combinations included a bleaching herbicide, suggesting synergy might be a class specific phenomenon. We also found that mesotrione-norflurazon and mesotrione-clethodim combinations remained synergistic against lettuce *(Lactuca sativa)*, but not tef *(Eragrostis tef)*. Our study shows that screening herbicide mixtures against *A. thaliana* is an efficient approach for finding rare herbicide synergies.

## Introduction

Since their implementation in agriculture in the 1940s, herbicides have improved crop productivity, but resistance now threatens those gains in yield. The first field case of herbicide resistance, triazine-resistant *Senecio vulgaris*, was documented in 1968^1^ and since then, cases of resistance have steadily risen with no signs of abatement.^2^ A common tactic to overcome resistant weeds is to switch between herbicides with different modes of action, but both experimental and computational modelling have shown that simple rotations do not delay the evolution of resistant weeds,^3, 4^ whereas more complex rotation patterns delay resistance only two-three times that from single herbicide use. A more efficient tactic is combining multiple herbicide modes of action in tank mixtures.^3^

Not all herbicides can or should be mixed with each other. Provided they are chemically compatible, the ideal herbicide mixtures contain active components with the same persistence and spectrum of controlled weeds, but a different mode of action.^5^ The activity of a herbicide mixture is not simply the sum of activity for the individual herbicides as one can affect uptake, translocation and metabolism of the other;^6^ meaning there are three types of herbicide interaction: additivity, antagonism or synergy.

Synergy is desirable when designing a herbicide mixture as it allows lower application rate or frequency of herbicide treatment, but finding a new synergy remains challenging. Sometimes, synergy can be hypothesised based on mechanistic assumptions, as it was done by Takano et al^7^ who predicted the synergy between glufosinate and protoporphyrinogen oxidase inhibitors and confirmed it experimentally, but generally synergies are not predictable. Despite frequent claims of synergy in patent literature, the number of peer-reviewed publications describing synergistic herbicide combinations is relatively low. Several hundred herbicide mixtures are described in patent and peer-reviewed literature; antagonism is 2-3 times more common than synergy, especially when herbicides from different chemical families are combined.^6, 8, 9^ Synergy was species dependent and seen more often for broadleaf than monocot weeds.^6, 9^ A synergistic herbicide mixture for one species can also be antagonistic or additive for another species ^8^. Thus, herbicide synergies appear to be rare and unpredictable.

The experimental data on herbicide mixtures is hard to compare, ranging in species tested with often a single synergy revealed in each publication. Most analyses are based on visual scoring of efficacy instead of absolute quantitative methods (e.g. shoot weight, leaf area measurement). This makes it hard to identify patterns in herbicide synergism, but certain groups of herbicides appear more likely to be synergistic than others. The literature on herbicide interactions from the last 20 years demonstrates the combination of the carotenoid synthesis inhibitor mesotrione and a PS II inhibitor atrazine to be one of the best-documented case of synergy, as it was independently confirmed by a number of research groups using both glasshouse and field studies (**Table 1**).

**Table 1.** Well documented cases of synergy. Mesotrione-atrazine and mesotrione-bromoxynil are two of the better documented and widely effective synergies reported in the literature.

| Synergy | Species | Fold | Reference |
| --- | --- | --- | --- |
| **mesotrione-atrazine** | *Abutilon theophrasti* | 1.5 to 6 | 16-18 |
|  | *Amaranthus palmeri* | 2 to 3 | 11, 16, 20 |
|  | *Amaranthus rudis* | 1.1 | 18 |
|  | *Ambrosia trifida* | 1.2 | 18 |
|  | *Chenopodium album* | 1.2 | 18 |
|  | *Ipomoea coccinea* | 1.5 to 4.3 | 10 |
|  | *Ipomoea hederacea* | 2.5 | 17 |
|  | *Raphanus raphanistrum* | 1.6 | 19 |
|  | *Setaria faberi* | 3.3 to 25 | 10 |
|  | *Xanthium strumarium* | 2.4 to 3.6 | 10 |
| **mesotrione-bromoxynil** | *Abutilon theophrasti* | 1.2 | 16 |
|  | *Amaranthus palmeri* | 2.5 | 16 |
|  | *Amaranthus rudis* | 1.2 | 34 |
|  | *Ambrosia trifida* | 1.2 | 34 |
|  | *Chenopodium album* | 1.2 | 34 |

The combination of atrazine and mesotrione was synergistic among a wide range of broadleaf and monocot weeds with the maximum synergy level reported to be 25-fold, when the herbicide mixture is applied to *S. faberi* after emergence.^10^ In addition to mesotrione and atrazine being a reliable synergy, combination of mesotrione and another PSII inhibitor bromoxynil was synergistic against *A. theophrasti* and *A. palmeri* which are same species for which the mesotrione-atrazine synergy was strongest (**Table 1**).

Similarly, mixing atrazine and an alternative inhibitor of carotenoid synthesis tembotrione was found to be synergistic against *A. palmeri.^11^* Atrazine was also found to be synergistic with clomazone, norflurazon and fosmidomycin,^10^ herbicides that similarly inhibit carotenoid biosynthesis, but via different mechanisms. This suggests a synergy found for a single herbicides might be indicative of a class-class synergy.

If synergies are a class-class phenomenon, it would be interesting to determine more systematically how common synergistic herbicide class pairs are and whether new ones can be found. Using controlled growth systems, a model plant and quantitative image analysis, we screened for synergy among the 276 pairwise combinations of 24 herbicides, each representing one mode of action. We found five reproducible synergies, namely atrazine-mesotrione, clethodim-mesotrione, atrazine-clomazone, paraquat-clomazone and norflurazon-mesotrione. All synergistic pairs included a bleaching herbicide (clomazone or mesotrione), and three combinations contained a reactive oxygen species activating component (atrazine or paraquat). Two combinations (mesotrione-atrazine and clomazone-atrazine) were already known, validating our ability to detect known, reliable synergistic interactions. Two newly discovered pairs (norflurazon-mesotrione and clethodim-mesotrione) worked both in *A. thaliana* and another dicot (lettuce, *Lactuca sativa*), whereas none of the synergies found were synergistic against a monocot (tef, *Eragrostis tef*). Our results show that a single-dose screening of herbicide combinations against *A. thaliana* could be extended to higher-throughput screening to find herbicidal synergies.

## Materials and Methods

### Materials

The majority of herbicidal active ingredients were purchased from Sigma Aldrich as analytical grade compounds: glyphosate (CAS 1071-83-6, catalogue number 45521), imazaquin (CAS 81335-37-7, catalogue number 37878), oxadiazon (CAS 19666-30-9, catalogue number 33382), norflurazon (CAS 27314-13-2, catalogue number 34364), mesotrione (CAS 104206-82-8, catalogue number 33855), clomazone (CAS 81777-89-1, catalogue number 46120), glyphosate (CAS 1071-83-6, catalogue number 45521), glufosinate-ammonium (CAS 77182-82-2, catalogue number 45520), chlorpropham (CAS 101-21-3, catalogue number 45393), dimethachlor (CAS 50563-36-5, catalogue number 45447), dinoseb (CAS 88-85-7, catalogue number 45453), ethofumesate (CAS 26225-79-6, catalogue number 45479), naptalam (CAS 132-66-1, catalogue number 33371), dazomet (CAS 533-74-4, catalogue number 45419), metam-sodium hydrate (CAS 137-42-8, catalogue number 45570), difenzoquat methyl (CAS 43222-48-6, catalogue number 34331). The compounds that were not available as analytical grade substance were purchased as the purest available substances from Sigma Aldrich: pelargonic acid (CAS 112-05-0, catalogue number N5502, purity 97%), paraquat dichloride (CAS 75365-73-0, catalogue number 856177, purity 98%), 2,4-dichlorophenoxyacetic acid (CAS 94-75-7, catalogue number D7299, purity 95%), dichlobenil (CAS 1194-65-6, catalogue number D57558, purity 97%), amitrol (CAS 61-82-5, catalogue number A8056, purity 95%). Atrazine (CAS 1912-24-9, catalogue number S334, purity 98%) and clethodim (CAS 99129-21-2, catalogue number O401, purity 90%) were purchased from AK Scientific. Asulam (CAS 3337-71-1, catalogue number A788750) was purchased from Sapphire Bioscience.

### Establishing sub-lethal herbicide dose

To screen for synergy requires sublethal doses of herbicides in pairs. Twenty-four herbicides were selected that represented roughly one per mode of action as well several in the unknown category (**Table 1**). Herbicide stocks were prepared as 12 mM solutions in dimethylsulfoxide (DMSO) except glyphosate, paraquat dichloride and glufosinate ammonium, which were dissolved in water. From this, a suite of herbicides stocks from 12 to 0.0015 mM were prepared by serial two-fold dilution of the 12 mM stock using DMSO. To prepare herbicide-containing medium, a 2.5 μL aliquot of a herbicide stock (or pure DMSO for negative control) was added to a well of a sterile transparent 96-well plate before it was filled with 250 μL of molten MS-agar medium consisting of 1% agar, 4 g/L Murashige-Skoog basal medium, 10 g/L glucose, 0.3% MES (w/v), pH 5.7. The final concentration of herbicide in agar wells ranged therefore from 120 μM to 0.015 μM. Surface sterilised *Arabidopsis thaliana* (Col-0) seeds were stratified in 0.1% agar for three days and 30-40 sown onto the surface of the agar medium. Once the surface had dried for ~15 min, plates were covered with lids and sealed with porous tape and transferred to a growth chamber under long-day illumination (16 h light/8 h dark, 136 μmol/m^2^/sec), 23°C and 60% relative humidity. After 14 days incubation, images were taken and plant growth was assessed visually. At some point across this concentration gradient, plants began to die and for each herbicide a second experiment was done using a finer gradient of concentrations to find the exact value for minimal efficient dose. Plant growth was quantified using the 14-day images using ImageJ (National Institutes of Health, 1.47v) with photos processed by ‘Threshold Colour’ at “Hue” −50-110, “Saturation” - 125-255 and “Brightness” 30-255 settings. These setting exclude background, yellow or white areas of plant leaves and retain only green pixels which correlate to healthy plants. Images were then converted into 8-bit format and then greyscale pixels were converted into red pixels that could be quantified by ImageJ. The green pixel count of plants treated by herbicides was normalised against the DMSO-only negative control to provide percentage inhibition. Inhibition values were plotted against herbicide concentration and the resulting dose-response curves were approximated by sigmoidal and hyperbolic curve models. The concentrations corresponding to the beginning of detectable growth inhibition were calculated by Hill’s equation and used as minimal effective concentration points in the subsequent synergy screening.

### Systematic synergy screening

To create a matrix for systematic synergy screening all possible pairwise combinations of herbicides were created. The 12 mM stocks were diluted with DMSO to a minimal effective concentration (MEC) for each herbicide, then equal volumes of individual herbicide stocks were mixed pairwise so every mixture had two herbicides each at 50% of their MEC. If the herbicide pair are additive, the inhibitory effect of the combination should not exceed 50% of inhibition, because none of single herbicides visibly inhibited plant growth at 50% of MEC. Herbicide pairs were then dissolved in agar as above. Each herbicide pair was tested in six replicates, whereas single herbicides were done in triplicate. After 14 days, incubation images were taken and plant growth quantified by ImageJ software package as above. The inhibition effect of each pair was averaged and compared to the expected effect based on dose response curves. To calculate the expected response, the Bliss independence model^12, 13^ was used:

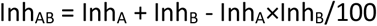

In this model, InhAB is expected percentage inhibition of a herbicides pair A and B, where each herbicide inhibits plant growth by InhA and InhB percentages respectively. We considered a combination synergistic if growth inhibition was two or more time the expected response calculated in assumption of pure additivity and three or more times the standard deviation.

### Generating isobolegrams for synergistic herbicides combinations

To confirm that herbicides synergies found in one dose screening are truly synergistic we generated isoboles. First, we generated dose-response curves for clethodim, atrazine, mesotrione and paraquat and approximated them by sigmoid dose response model. Then, we graphically determined the concentrations corresponding to 50% of growth inhibition (EC_50_) and used these values as starting points to create a grid of herbicide mixtures containing different concentrations of synergistic herbicides at different ratios. To do so we prepared stocks that contained each herbicide at 90 - 10% of EC_50_ and made all possible pairwise combinations. These were dissolved in agar as described above. After 14 days incubation, images were taken and plant growth quantified by ImageJ software package as above. We then chose the data points where inhibition was 50±10% and plotted them on Cartesian plot, where the concentration of one herbicide was used as an abscissa, and the concentration of another herbicide was used as an ordinate. The experiments were repeated three times, and if the concentration of herbicides resulting in 50% inhibition varied between replicates, then the herbicide concentrations were averaged and plotted with error bars representing standard deviation.

### Herbicidal activity assay on soil

To confirm whether synergies discovered on agar plates were relevant, the activity of individual herbicides and herbicide mixtures was compared on soil according to the method from Corral et al. ^14^. Roughly 30 *Arabidopsis thaliana* Col-0, or 20 tef *(Eragrostis tef)* or 10 lettuce *(Lactuca sativa)* seeds were sown in 63 x 63 x 59 mm pots consisting of Irish peat that was pre-wet before sowing. Seeds were stratified for 3 days *(A. thaliana)*, 5 days *(L. sativa)* or 7 days *(E. tef)* in the dark at 4°C and grown in a chamber at 23°C, with 60% relative humidity and in a 16 h light / 8 h dark photoperiod. The individual herbicides and herbicide mixtures were prepared as stocks in DMSO and diluted in water prior to treatment, so the final concentration of DMSO was 2%. The polyether modified polysiloxane based surfactant Brushwet (SST Australia) was added to a final concentration of 0.02% (v/v). The 2% DMSO solution in water contained 0.02% (v/v) of Brushwet was used as a negative control. To establish the right dose range for synergistic herbicides, a series of solutions containing 400 mg/L - 0.001 mg/L of herbicide was prepared by serial two-fold dilutions and with 500 μL of herbicidal solutions applied to seeds (pre-emergence treatment) or seedlings (post-emergence treatment) by a pipette (*A. thaliana* and lettuce) or by spraying (tef). Pre-emergence treatments were given as trays were moved into their first long day, whereas post-emergence treatments were done three and six days after germination. The minimal concentration corresponding to the visible growth inhibition was used as a starting point for preparing herbicide mixtures. Seedlings were grown for 16 days after transfer to the growth room before images were taken, and plant growth quantified. For *A. thaliana*, plant growth was quantified by measuring green pixels as described above, for *E. tef* and *L. sativa* the growth was quantified by measuring fresh weight. To quantify fresh weight of the plants, they were cut at the height of 3-5 mm from soil using sharp scissors and weighed immediately.

### Assessment of synergistic response

To assess if a herbicidal pair was synergistic, individual herbicides and their mixtures were tested against model plants in eight replicates. The inhibitions percent were normalised against control plants treated with 2% DMSO solution in water contained 0.02% (v/v) of Brushwet and averaged. The inhibition percentage for individual herbicides were used to calculate the expected inhibition percentage of herbicide mixture using the Bliss independence model.^12, 13^ The observed inhibition percentage for herbicidal mixtures were compared to the expected inhibition percentage values using one sample t-test. The mixture was considered as synergistic if the observed response exceeded the expected and p-value was smaller than 0.05. If the expected response significantly exceeded the observed inhibition percentage, the mixture was considered as antagonistic. If there were no significant difference between expected and observed inhibition percentage, the mixture was considered as additive.

## Results

To find which herbicides were synergistic we created a matrix of 24 herbicides, where 20 herbicides represented one mode of action each, and four herbicides had an unknown mode of action (**Table 2**). The herbicides chosen for screening are widely used and included three molecules (clomazone, mesotrione and atrazine) for which well-documented cases of synergy are known as positive controls.^10, 11, 15–20^

**Table 2.** Herbicides used for synergy screening. Modes of action and herbicides named with WSSA recommendations. Classifications include the 2020 global classification and, in brackets, the legacy Herbicide Resistance Action Committee classification. In the far-left column is the letter code used in this work (**Figure 1**).

| Code | Herbicide common name | Class | Mode of action |
| --- | --- | --- | --- |
| A | clethodim | 1 (A) | Inhibits acetyl CoA carboxylase |
| B | imazaquin | 2 (B) | Inhibits acetolactate synthase |
| C | atrazine | 5 (C1,2) | Inhibits photosynthesis at PS II |
| D | paraquat dichloride | 22 (D) | PS I electron diverter |
| E | oxadiazon | 14 (E) | Inhibits protoporphyrinogen oxidase |
| F | norflurazon | 12 (F1) | Inhibits phytoene desaturase |
| G | mesotrione | 27 (F2) | Inhibits 4-hydroxyphenyl-pyruvate dioxygenase |
| H | amitrole | 34 (F3) | Inhibits lycopene cyclase |
| I | clomazone | 13 (F4) | Inhibits 1-deoxy-D-xylulose 5-phosphate synthase |
| J | glyphosate | 9 (G) | Inhibits 5-enolpyruvyl-shikimate-3-phosphate synthase |
| K | glufosinate ammonium | 10 (H) | Inhibits glutamine synthetase |
| L | asulam | 18 (I) | Inhibits 7,8-dihydro-preroate synthetase |
| M | oryzalin | 3 (K1) | Inhibits microtubule assembly |
| N | chlorpropham | 23 (K2) | Inhibits microtubule organization |
| O | dimethachlor | 15 (K3) | Inhibits very long chain fatty acid synthesis |
| P | dichlobenil | 29 (L) | Inhibits cell wall synthesis |
| Q | dinoseb | 24 (M) | Uncoupler of oxidative phosphorylation |
| R | ethofumesate | 15 (K3) | Inhibits very long chain fatty acid synthesis |
| S | 2,4-dichlorophenoxyacetic acid | 4 (O) | Synthetic auxin |
| T | naptalam | 19 (P) | Inhibits indoleacetic acid transport |
| U | dazomet | 0 (Z) | Unknown mode of action |
| V | metham sodium hydrate | 0 (Z) | Unknown mode of action |
| W | difenzoquat methyl sulfate | 0 (Z) | Unknown mode of action |
| X | pelargonic acid | 0 (Z) | Unknown mode of action |

To detect synergistic herbicide combinations, the 276 possible pairwise mixtures of 24 herbicides were made and screened using *Arabidopsis thaliana* grown on agar plates. For each possible combinations (276), a single sub-lethal concentration for each herbicide was used. Both herbicides were used at 50% of their minimal effective concentration (MEC), so inhibition should not be visible if pairs were only additive. The concentrations were chosen such that even ~1.5-fold synergy would be easily observable as lethal or growth inhibition across a ‘green background’ of additivity. Individual herbicides at MEC and 50% of MEC were used to ensure the selected doses were indeed within the sub-lethal range.

Growth inhibition was quantified by measuring green leaf area and normalised against data from plants grown on media with no herbicide. Then the data were averaged and visualised as heat map (**Fig. 1A-B**).

**Fig. 1.**
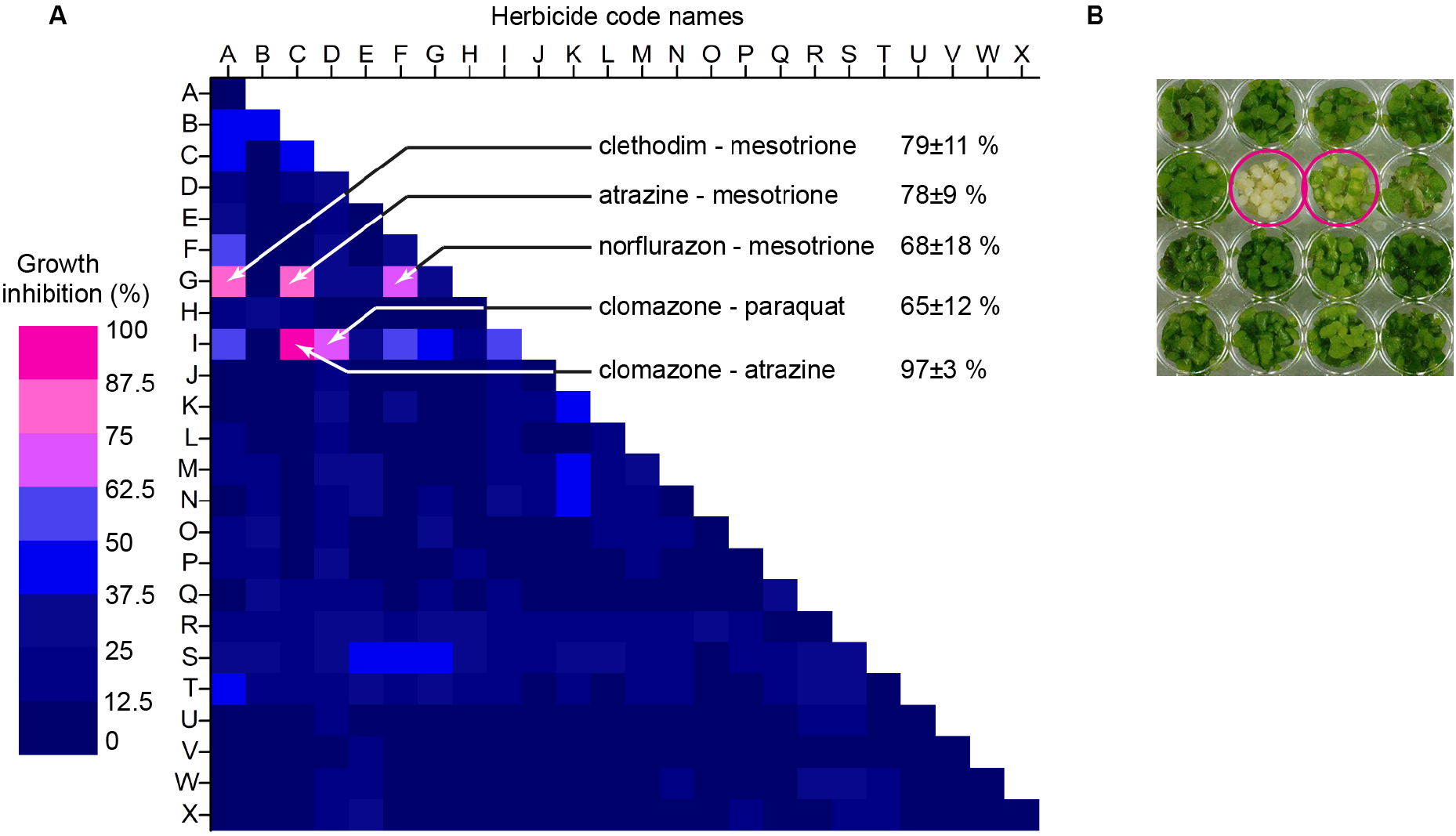
Five of 276 herbicide pairs were synergistic and all five included either mesotrione or clomazone. (A) Heat map summarising effect of herbicide pairs on growth of *A. thaliana*. Each herbicide mixture was replicated in independent experiments six times and single herbicides (24, e.g. A+A) were replicated three times. Herbicide identity (A-X) are listed in **Table 1**. (B) A typical image of agar plate used for screening. Synergistic combinations are marked by magenta circles.

Only five of 276 combinations were synergistic, namely atrazine-mesotrione, clethodim-mesotrione, norflurazon-mesotrione, atrazine-clomazone, and paraquat-clomazone. Three herbicides stood out in these synergies; three of the pairs included mesotrione, two included clomazone and two included atrazine. Two combinations (mesotrione-atrazine and clomazone-atrazine)^10, 11, 15–20^ were previously reported in the literature, which showed that one-dose screening was sensitive enough to detect these known synergistic mixtures.

To confirm the synergies identified using herbicide pairs, each at sub-lethal doses, we generated isobolegrams for each pair where the concentrations of each ranged. The approach involved plotting the concentration that causes 50% of growth inhibition (EC_50_) for one herbicide as a point on the X axis of a Cartesian plane, and the EC_50_ of the second herbicide as a point on the Y axis. A straight line (isobole) drawn between these points will represent the concentration of mixtures of this pair that should give 50% growth inhibition if there were no interaction between the pair (i.e. additivity).^21, 22^ If an herbicide pair is synergistic, the dose needed for 50% inhibition as a pair will be lower, lying below the line for additivity. If herbicides are antagonistic, the doses needed for 50% inhibition as a pair will be higher and so above the line for additivity.

To find combinations that give 50% inhibition we established EC_50_ values for synergistic herbicides using dose response curves approximated by sigmoid dose response model (**Fig. 2**). Then we prepared a series of mixtures where concentration of one herbicide was constant, and concentration of another herbicide varied from 10% of EC_50_ to the dose that lies on additive line. In total, for each herbicide pair we prepared 39 mixtures covering the area under the additivity line with an even grid of herbicide concentrations (**Fig. 3**).

**Fig. 2.**
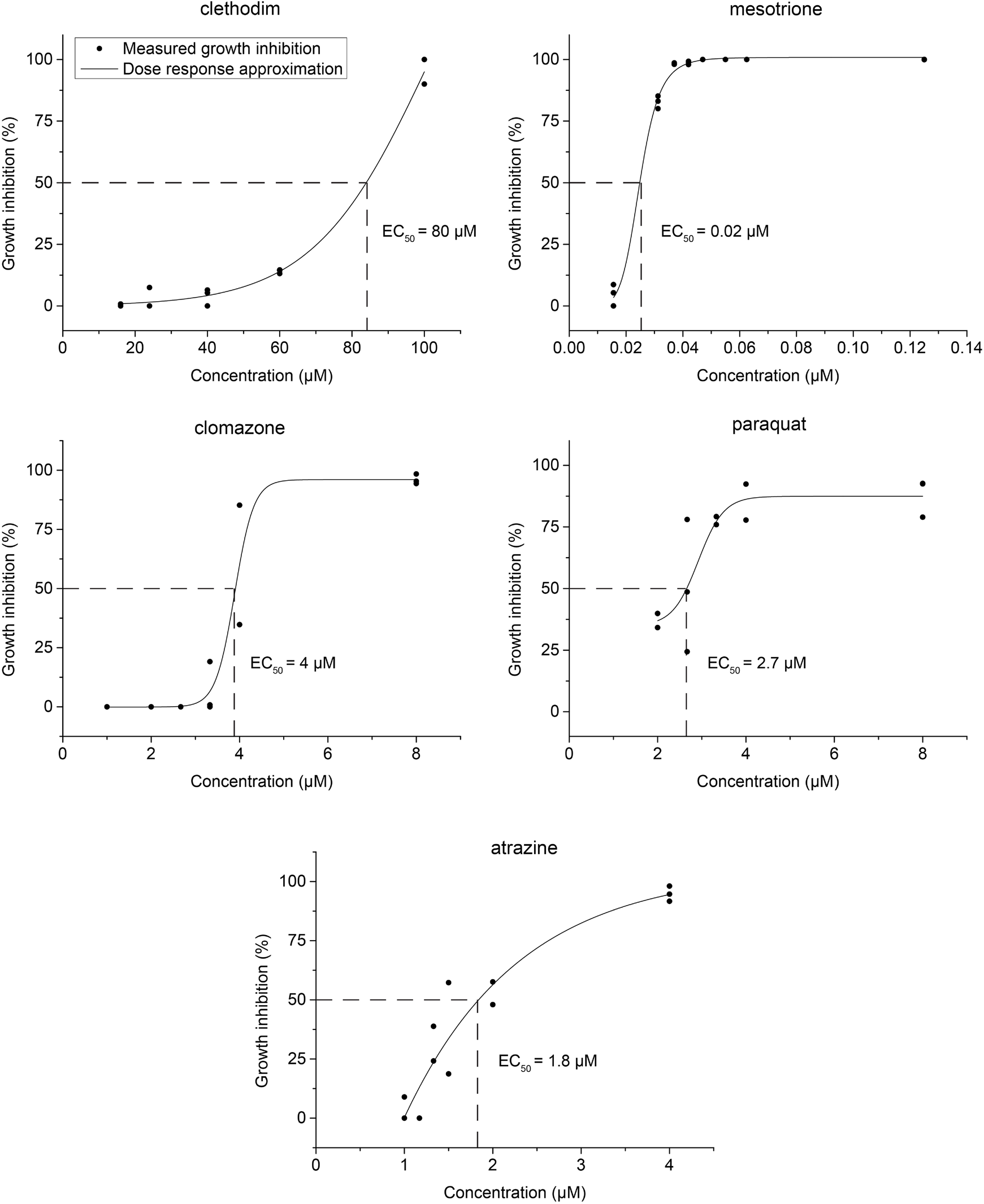
Mesotrione was the most active among synergistic herbicides. Dose-response curves were generated by agar plate germination assay. Clethodim was still active against *A. thaliana* despite being used for its strong effects against grasses relative to broadleaf plants.

**Fig. 3.**
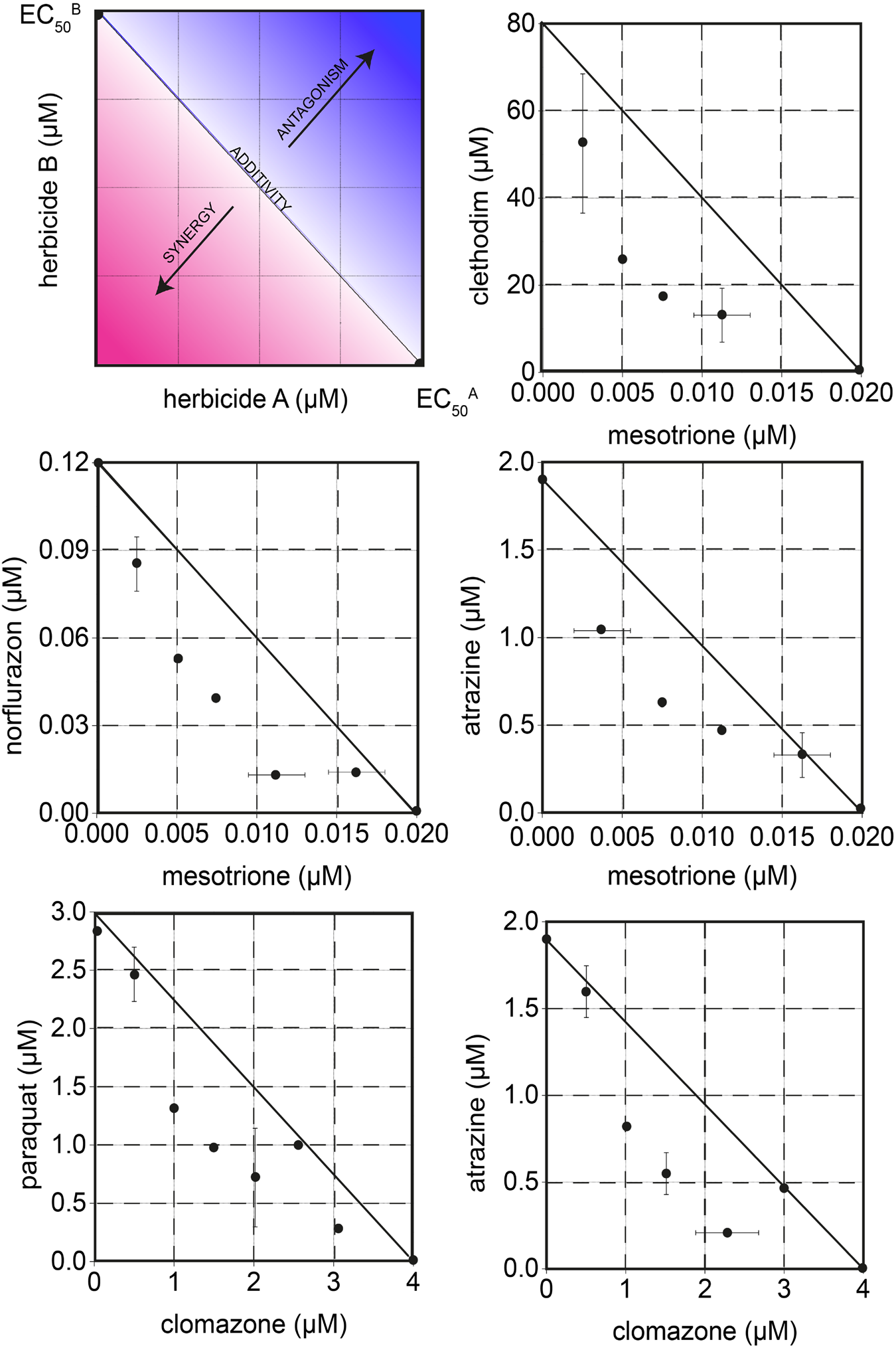
Isobolegrams of all five herbicide pairs show synergism in the dose range near EC_50_. The line (isobole) between the EC_50_ of two herbicides represents additivity. Mixtures below these concentrations that give 50% inhibition are synergistic and fall below the line. If higher concentrations are needed in mixtures to achieve 50% inhibition this represents antagonism and values fall above the line. Error bars represent standard deviation.

The isobolegrams show that clethodim-mesotrione (**Fig. 3**) was synergistic at all the clethodim to mesotrione ratios tested. By contrast, mixtures of mesotrione with atrazine (**Fig. 3**) or norflurazon (**Fig. 3**) were close to additive at ratios skewed highly toward the concentration that cause 50% of growth inhibition of one of the pair. Similarly, mixtures of clomazone and paraquat (**Fig. 3**) or atrazine (**Fig. 3**) were synergistic at most ratios, but additive at ratios skewed highly toward one herbicide.

Although the initial sub-lethal dose paired screen (**Fig. 1**) and isobolegrams (**Fig. 3**) are consistent, both are done on agar plates. To determine if these combinations identified remain synergistic in more natural growth conditions, we tested the same five herbicide pairs against soil-grown *Arabidopsis thaliana*. The herbicidal mixtures or individual herbicides were applied directly to seeds on their first day (pre-emergence treatment) or to seedlings on the third and sixth day after germination (post-emergence treatment). Some of the tested herbicides had different efficiency depending on the stage of application, therefore we adjusted doses accordingly to avoid full inhibition of plants growth by single compounds. Growth inhibition was again quantified by green leaf area and compared to plants not treated by herbicides (**Fig. 4–7**).

**Fig. 4.**
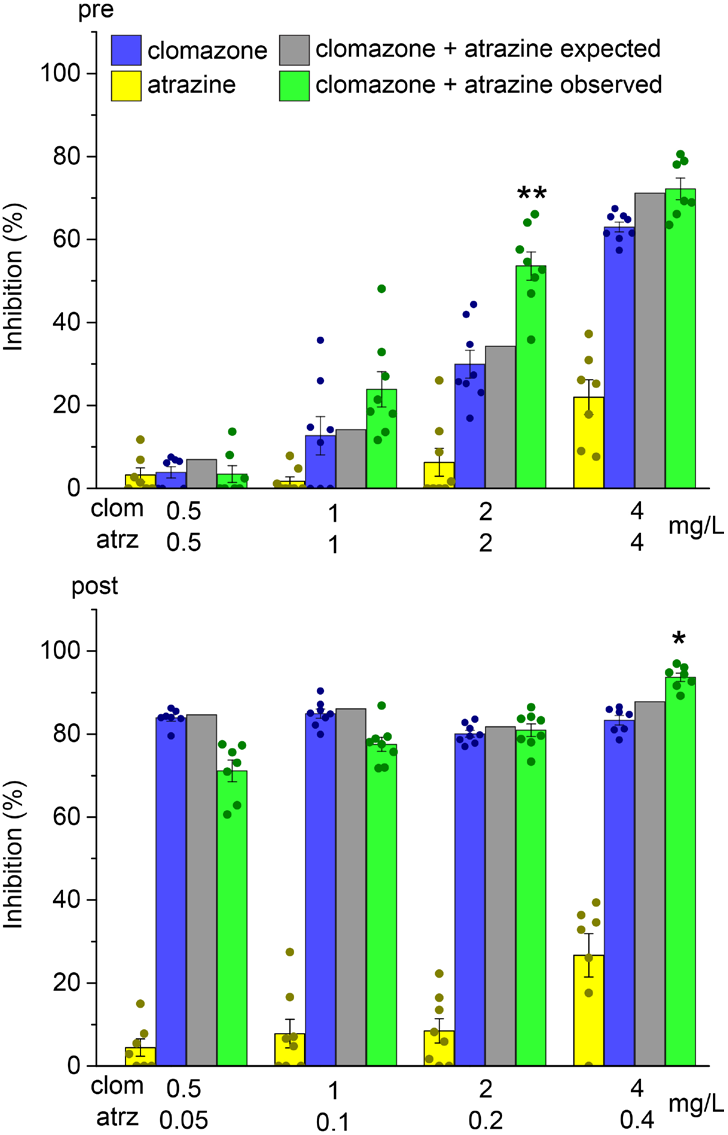
Clomazone and atrazine are more synergistic pre-emergence in mid-dose range. Clomazone and atrazine were moderately synergistic against *A. thaliana* when applied pre-emergence at 2 mg/L of clomazone and 2 mg/L atrazine. Some synergy was also seen post-emergence at the highest dose-range doses. Error bars represent standard error (SE). Asterisks denote the significance of difference from expected response for additivity, * p < 0.05, ** p < 0.01, *** p < 0.001.

**Fig. 5.**
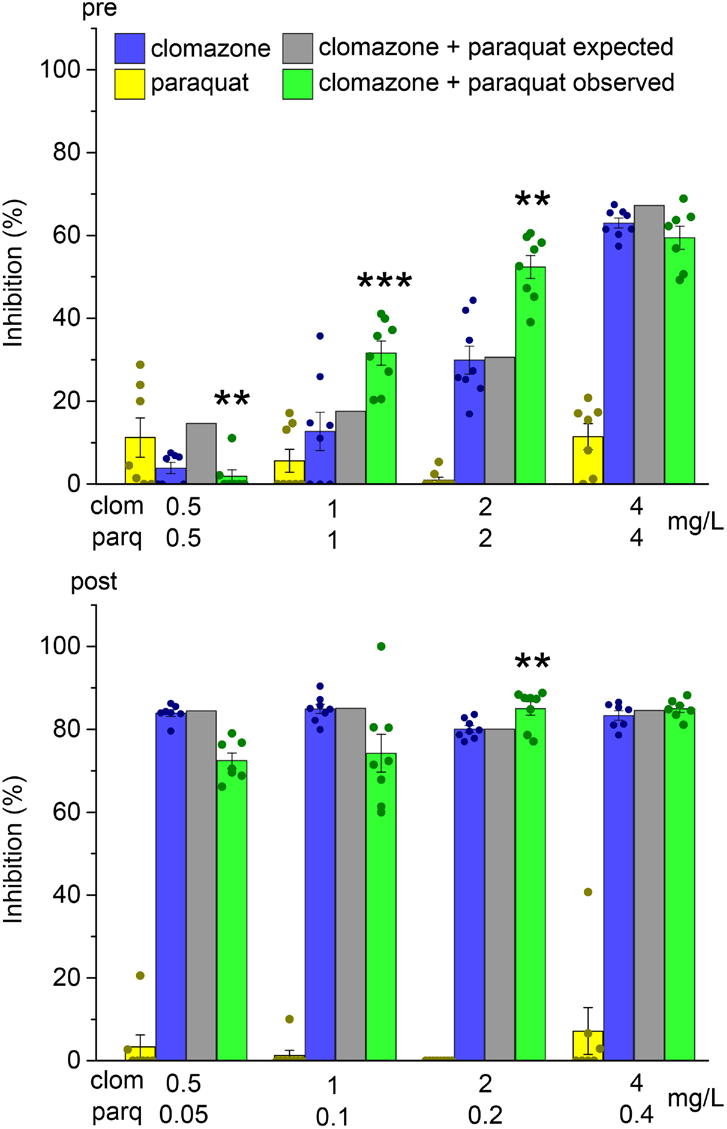
The clomazone-paraquat synergy is more synergistic pre-emergence in mid-dose range. Clomazone and paraquat were moderately synergistic against *A. thaliana* pre-emergence at doses of paraquat and clomazone between 1 and 2 mg/L, and mostly additive post-emergence. Error bars represent SE. Asterisks denote the significance of differences from that expected for additivity: * p < 0.05, ** p < 0.01, *** p < 0.001.

**Fig. 6.**
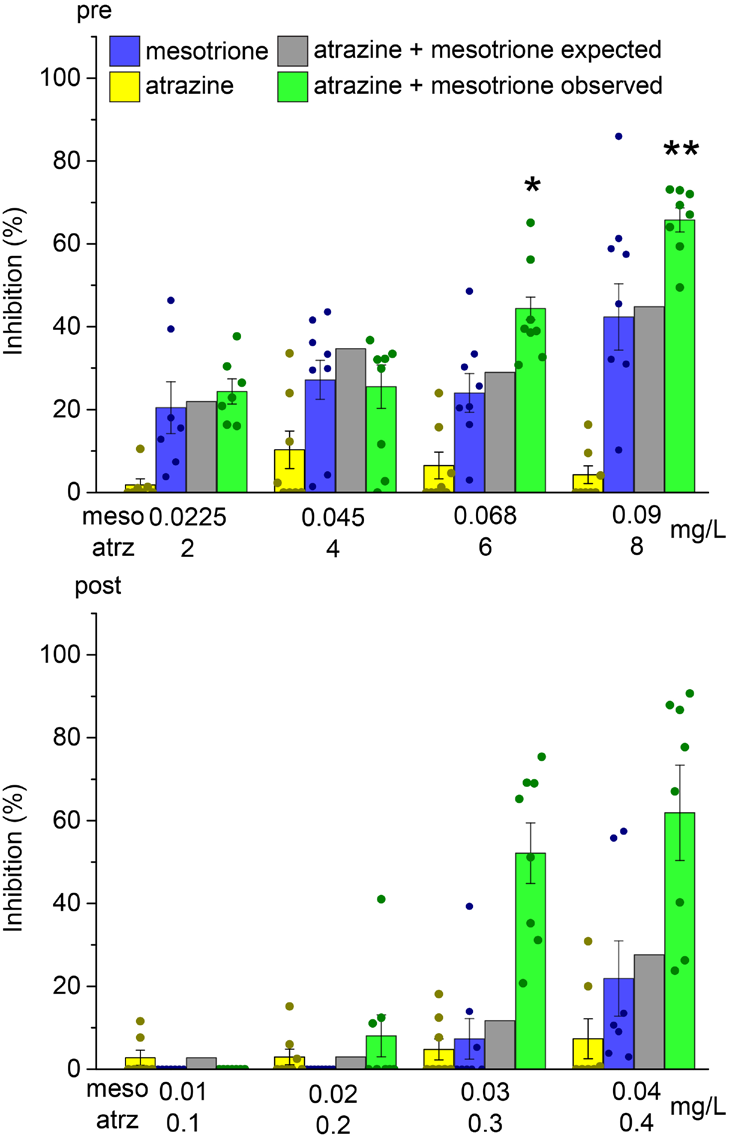
Mesotrione-atrazine are more synergistic at higher rates |. The biggest synergy against *A. thaliana* was 1.3-fold and observed for mixture of 0.09 mg/L of mesotrione and 8 mg/L of atrazine applied pre-emergently. Error bars represent SE. Asterisks denote the significant difference with expected response for pure additivity: * p < 0.05, ** p < 0.01, *** p < 0.001.

**Fig. 7.**
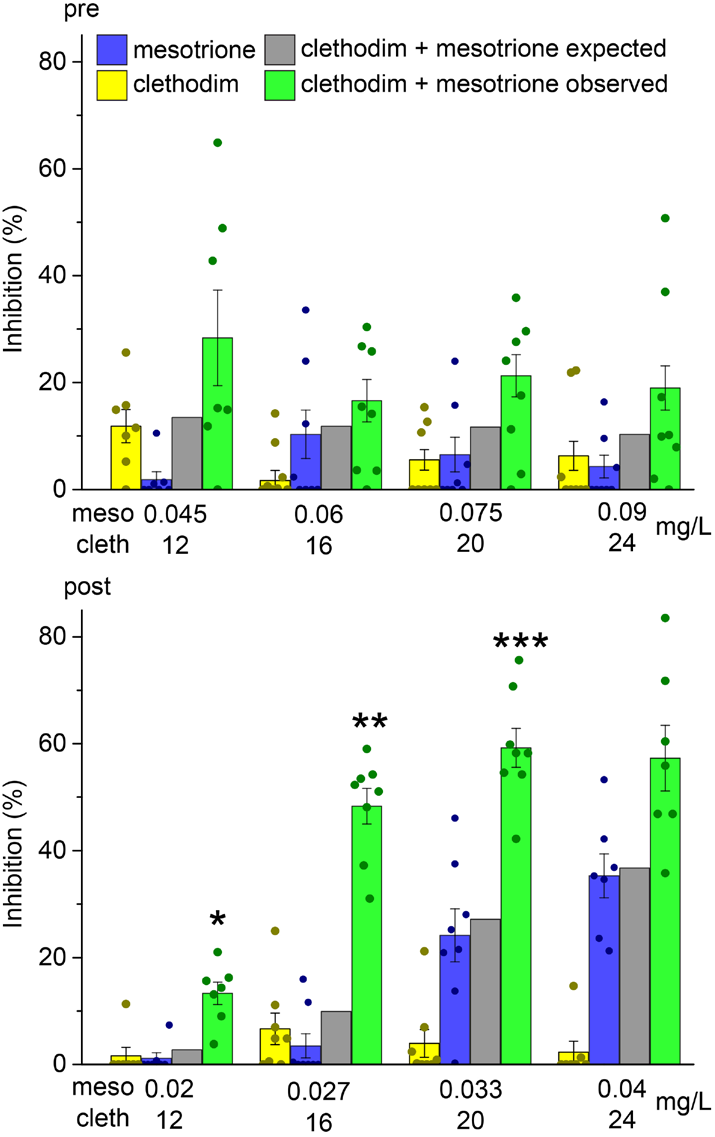
Mesotrione and clethodim were synergistic post-emergence, except at the highest dose. The synergy against *A. thaliana* was up to 4-fold for 0.027 mg/L of mesotrione and 16 mg/L of clethodim. Error bars represent SE. Asterisks denote the significance of differences from expected for additivity * p < 0.05, ** p < 0.01, *** p < 0.001.

In general, the synergy for the pairs remained, but was less marked on soil-grown plants. The clomazone-atrazine combination was moderately synergistic at the highest application rate post-emergence; this mixture inhibited plant growth by ~8% more than the expected effect calculated from the *A. thaliana* response to the individual herbicides at the corresponding concentrations. The synergistic effect was higher for pre-emergence application and reached ~1.4-fold increase in herbicide efficiency in the mid-dose range (**Fig. 4**).

For the clomazone-paraquat combination, moderate synergy was observed in the mid-range dose applied pre-emergence, whereas the lowest dose was antagonistic. When applied post emergence there was a slight synergy in the mid-dose range, and additivity at other application rates (**Fig. 5**).

Mesotrione and atrazine were ~1.3-fold synergistic at high doses pre-emergence, but additive at lower rates. Similarly, mesotrione-atrazine mixture was additive at lower doses and synergistic at higher application rates when applied post-emergence (**Fig. 6**).

Unlike the clomazone-atrazine and clomazone-paraquat mixtures we discussed above, that were synergistic regardless of application stage the mixture of clethodim and mesotrione was synergistic only post-emergence (**Fig. 7**). The synergistic response was observed up to the maximal application rate and varied from 2- to 4-fold depending on dose.

The mixture of mesotrione and norflurazon was also sensitive to the application stage: mesotrione was synergistic with norflurazon only when applied pre-emergence at the highest dose (**Fig. 8**).

**Fig. 8.**
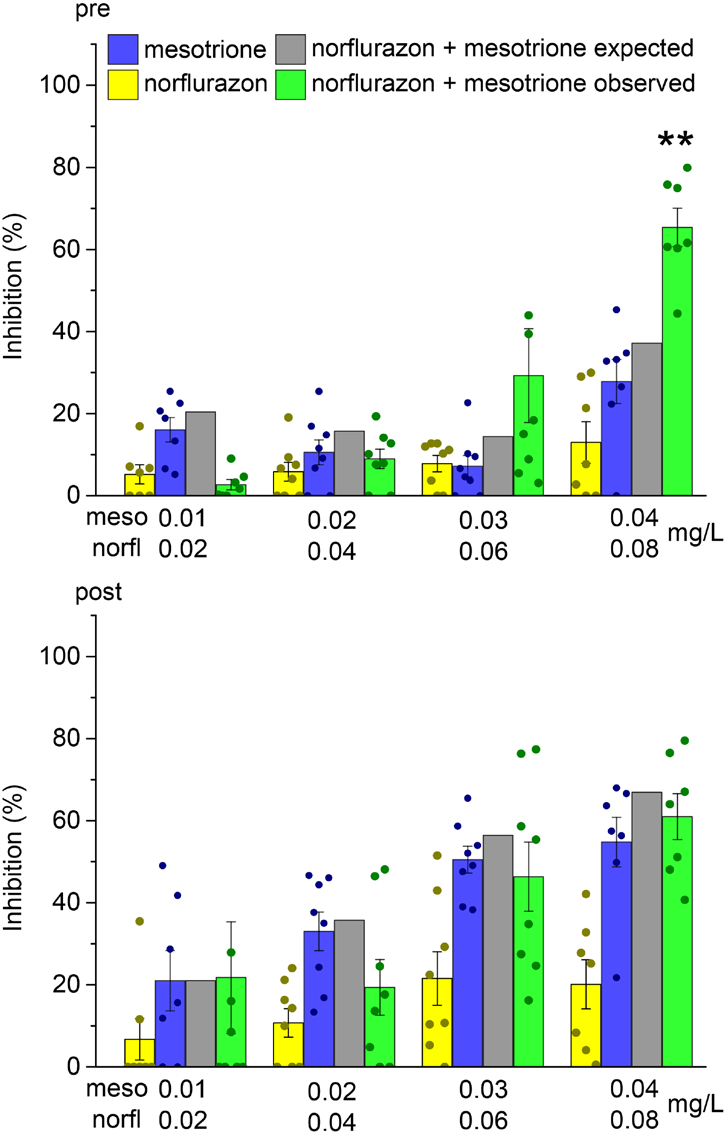
Mesotrione and norflurazon were synergistic pre-emergence, only at the highest dose. Synergism was observed for mixture of 0.04 mg/L of norflurazon and 0.04 mg/L of mesotrione when applied pre-emergence against *A. thaliana*. Error bars represent SE. Asterisks denote the significance difference from expected response for additivity * p < 0.05, ** p < 0.01, *** p < 0.001.

Overall, synergies found on agar plates worked on soil, albeit more weakly. To test whether the herbicide combinations that we found may have a broader applications we sought to determine whether the synergistic combinations worked in species other than *A. thaliana*. We tested the discovered synergistic combinations (clomazone-paraquat, clethodim-mesotrione and norflurazon-mesotrione) against lettuce *(Lactuca sativa)*. We chose lettuce as it germinates efficiently, has small seeds that are conveniently available, is easy to handle, and it is not a close evolutionary relative of *A. thaliana*. The herbicide mixtures and individual herbicides were applied to soil-grown lettuce in the similar manner to *A. thaliana;* the herbicidal mixtures or individual herbicides were applied directly to seeds on their first day (pre-emergence treatment) or to seedlings on the second and fifth day after germination (post-emergence treatment). Growth inhibition was quantified by measuring fresh weight of the above soil part of the plants. As lettuce is not as planar as *A. thaliana* quantification by green pixels count is a less accurate measure of growth. We found that all combinations remained synergistic in lettuce (**Fig. 9–11**), albeit to a lesser extent than *A. thaliana*.

**Fig. 9.**
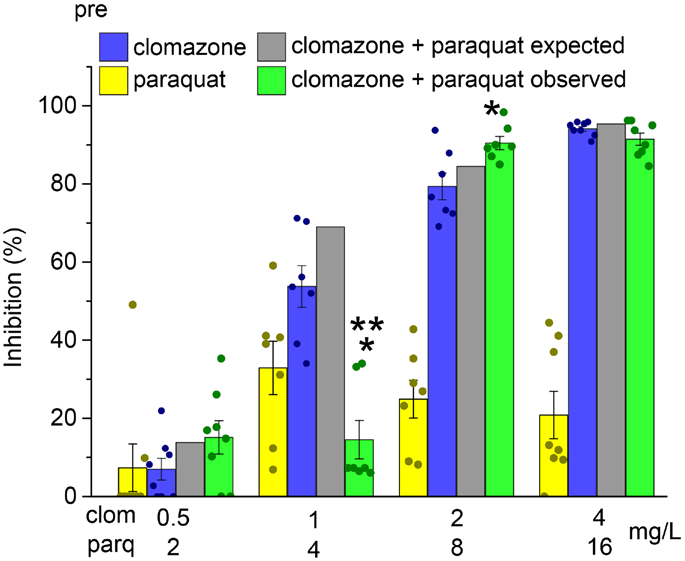
The clomazone-paraquat synergy depends on the dose. Clomazone and paraquat were slightly synergistic against lettuce *(L. sativa)* pre-emergence at doses of paraquat and clomazone 2 and 8 mg/L, and antagonistic at lower and higher doses. Error bars represent SE. Asterisks denote the significance of differences from that expected for additivity: * p < 0.05, ** p < 0.01, *** p < 0.001.

**Fig. 10.**
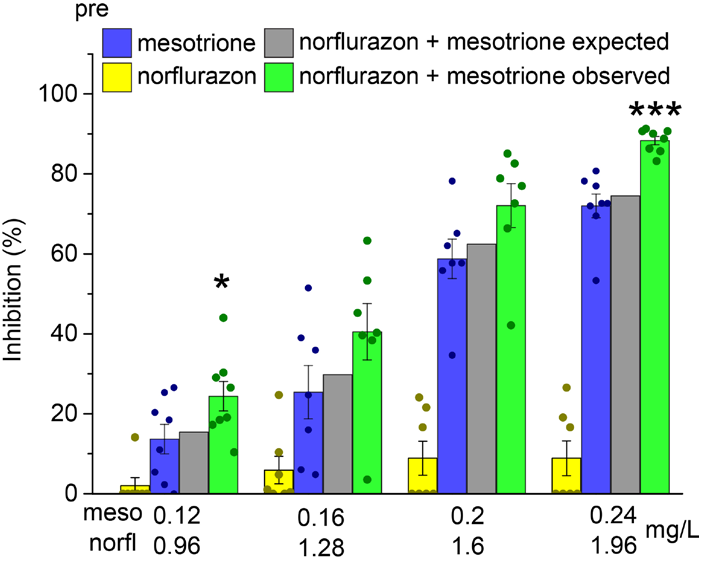
Mesotrione and norflurazon were synergistic at all doses. The highest synergism was observed for mixture of 0.96 mg/L of norflurazon and 0.12 mg/L of mesotrione when applied pre-emergence against *L. sativa*. Error bars represent SE. Asterisks denote the significance difference from expected response for additivity * p < 0.05, ** p < 0.01, *** p < 0.001.

**Fig. 11.**
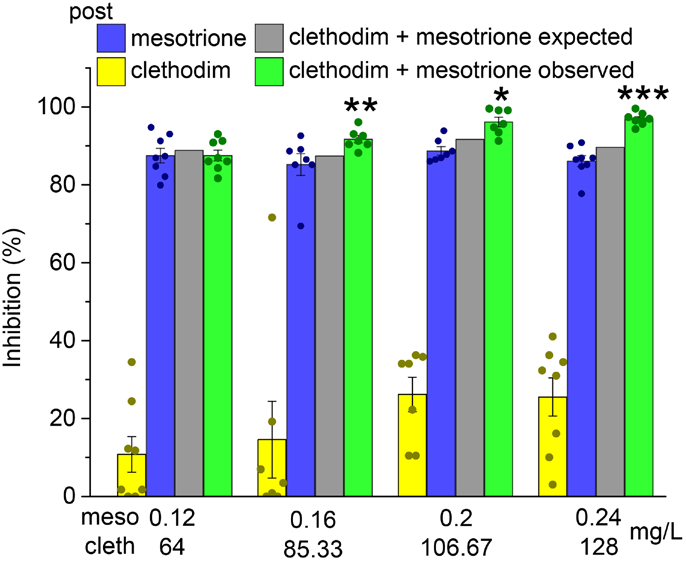
Mesotrione and clethodim were synergistic post-emergence, when applied at higher doses. The synergy against *L. sativa* was small (8-10%) but detectable starting from the mid low dose range. Error bars represent SE. Asterisks denote the significance of differences from expected for additivity * p < 0.05, ** p < 0.01, *** p < 0.001.

As was seen for *A. thaliana*, the combination of paraquat and clomazone had a complex dependence of synergy on dose in *L. sativa* too: the mixture was additive at the lowest dose, strongly antagonistic when herbicide dose was doubled and we observed a ~1.1-fold synergy when the dose increased again and was a slight antagonism with further increase of the application rate (**Fig. 9**).

In contrast the mesotrione-norflurazon mixture was synergistic at all tested doses against *L. sativa*. The degree of synergy varied between ~1.3 fold at the lowest dose and ~1.2 fold at the highest of increased herbicidal effect for the mixture as compared to expected response and was largest in the low and high-dose range (**Fig. 10**).

Clethodim-mesotrione combinations was more synergistic at higher doses when applied to in *L. sativa* (**Fig. 11**), unlike for *A. thaliana* where it was more synergistic at lower doses.

To further extend our observation from *A. thaliana* we tested mesotrione-clethodim, paraquat-clomazone and mesotrione-norflurazon mixtures against a monocot species tef *(Eragrostis tef)*. None of the combinations were synergistic against tef: mesotrione-norflurazon and mesotrione-clethodim combinations were antagonistic, whereas paraquat-clomazone mixture was additive (**Supp Fig. 1–3**). These findings were consistent with previous observations that synergies are more common in broadleaf species than in monocots.^6, 9^

## Discussion

We used a single-dose screen of herbicide mixtures against model plant *A. thaliana* as a tool to discover synergistic combinations. We created a screen of 276 pairwise combinations from 24 herbicides with different modes of action. Although the herbicide doses corresponding to ~50% of inhibition are considered the most suitable for synergy detection^13, 23^ we used sub-lethal concentrations because in this case the difference between observed and expected inhibition will be larger, so the variation between replicates and errors of quantification should not affect the detection of synergy. We used a germination assay on agar plates because it allowed us to screen all possible combinations simultaneously, under controlled conditions and analyse the growth quantitatively. To avoid confounding herbicide interactions with interactions caused by chemicals in the herbicide formulations we used pure, herbicide actives alone in agar and with only a surfactant (polyether modified polysiloxane) and carrier (DMSO) in follow-up soil treatments. Commercial herbicide formulations typically contain surfactants, solvents, antifoaming agents and buffers^24^ so combining formulations as well as herbicide actives would have created more complexity. Surfactant efficiency also is known to depend on herbicide chemistry and species.^25^

The single-dose screen revealed five herbicide combinations namely clomazone-atrazine, mesotrione atrazine, mesotrione-norflurazon, mesotrione-clethodim and paraquat-clomazone. These were confirmed to be truly synergistic by creating isobole plots with data from plants grown under the same conditions. All combinations were later tested against soil-grown *A. thaliana* and remained synergistic. Among these synergies, paraquat-clomazone might look confusing at the first glance as it worked best when applied pre-emergence, although paraquat is typically a contact herbicide. However, it was shown, that being sprayed on peat soil, paraquat does not absorb fully^26^ and forms a thin layer of active compound on soil surface,^27^ so the seeds sown close to the surface are exposed to paraquat and demonstrate reduced growth.^26–28^ In our experimental design, *A. thaliana* seeds were sown on the surface of Irish peat, thus creating the conditions where pre-emergence activity of paraquat is maximal, so the observed synergy effect for pre-emergence application of paraquat-clomazone mixture is not surprising. Another unexpected result was the synergy between clethodim and mesotrione, because clethodim more strongly affects grass species^29^, so *A. thaliana* should be clethodim-tolerant. However, we observed clethodim to be moderately active against *A. thaliana,^30^* therefore synergy between mesotrione and clethodim was possible. Thus, we can conclude that single-dose screen on agar plates is sufficient to detect synergistic herbicide combinations; specifically it can distinguish between additivity and synergy.

We confirmed that the method is sensitive enough to reliably find synergies by comparing our results with recent synergy claims in peer-reviewed literature. Analysis of the synergy literature published in the last two decades showed that the 24 herbicides chosen for screening can potentially form five synergistic combinations: mesotrione-atrazine,^10, 11, 16–20^ clomazone-atrazine,^15^ paraquat-atrazine,^31^ norflurazon-atrazine^10^ and glufosinate-2,4 D.^32^ Two combinations (mesotrione-atrazine, clomazone-atrazine) were rediscovered in our screen, but three were not.

The simplest explanation for this is that synergies can be species-specific or concentration-dependent as we saw for clomazone-paraquat that was only synergistic at one, mid-range concentration and did not extend from *A. thaliana* to *L. sativa*.

The glufosinate-2,4-D synergy which we did not see in *A. thaliana*, was synergistic in *Commelina communis* and *Echinochloa colona*, but additive in *Ambrosia trifida*^32, 33^ The paraquat-atrazine synergy^31^ was only reported for monocot weed species *Echinochloa colona* and *Chloris virgata*. Other factors that could complicate synergies are that many studies use formulated herbicides, not pure actives. Others used alternative carriers and surfactants, which can be important. Work made by Liu using three different species (monocot *Triticum aestivum* and two dicots *Chenopodium album* and *Vicia faba)*, two different herbicides (2,4-D and glyphosate) and 22 different surfactants showed that uptake of herbicide depends on both, herbicidal molecule and surfactant structure, as well as plant species,^25^ therefore synergy might depend on which surfactant and weed species are used. Armel et al.^10^ working with *Xanthium strumarium, Setaria faberi* and *Ipomoea coccinea* used mixture of acetone, glycerol and water as a carrier and a polyoxyethylene surfactant when finding a clomazone-norflurazon synergy whereas we used *A. thaliana*, DMSO and polysiloxane respectively as species, carrier and surfactant.

Nevertheless, a single-dose screen with *A. thaliana* was sensitive enough to re-discover well-documented synergies with broad species sensitivity. To check if the three combinations newly discovered using *A. thaliana* remained synergistic in other species we tested them against broadleaf (lettuce) and one monocot (tef) model organisms. We found that two pairs (norflurazon-mesotrione and clethodim-mesotrione) remained synergistic from *A. thaliana* to lettuce. For the combination of paraquat and clomazone all types of interaction (synergism, antagonism and additivity) were observed depending on concentration, so this pair cannot be classified simply as synergistic. By contrast, none of the tested combinations were synergistic in tef, consistent with many synergies being species-specific as well as being more common to dicots than monocots.^9^ As synergies found in *A. thaliana* reproduced well in another dicot, but not a monocot species suggests initial screening for synergies that work against monocot weeds should be done with a monocot species, like tef.

## Conclusion

We developed a reliable tool for discovering synergistic herbicide combinations. Using a quantitative method, we detected previously identified reliable synergies and found new ones that have potential to act across a broad range of dicot weed species. Interestingly, all synergistic combinations detected in our screen contained at least one carotenoid biosynthesis inhibitor (clomazone, norflurazon or mesotrione). This observation suggests a more detailed study of the mechanism of interaction between bleaching herbicides and other classes would be beneficial to finding more herbicide synergies.

## Acknowledgements

K. V. S. was supported by a Research Training Program Stipend and UWA Safety-Net Top-Up Scholarship. This work was funded in part by Nexgen Plants and in part by an Australian Research Council Discovery Project (DP190101048) to J. S. M. The authors thank Philippe Hervé and Bruce Lee from Nexgen Plants for the useful comments on the manuscript. The authors are grateful to and Jens Lerchl and his colleagues from BASF, and Hudson Takano from Corteva Agriscience for helpful input based on the earlier preprint version of the manuscript held on *bioRxiv*.

## Conflict of Interest Declaration

There are no conflicts to declare

## Supplementary figures

**Supplementary Fig. 1.**
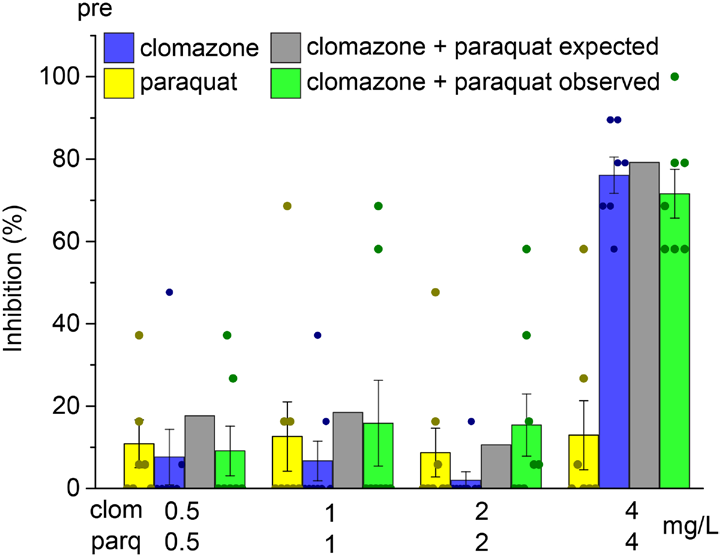
The clomazone-paraquat was additive in tef. Clomazone and paraquat were additive at all doses when applied pre-emergence. Error bars represent SE. Asterisks denote the significance of differences from that expected for additivity: * p < 0.05, ** p < 0.01, *** p < 0.001.

**Supplementary Fig. 2.**
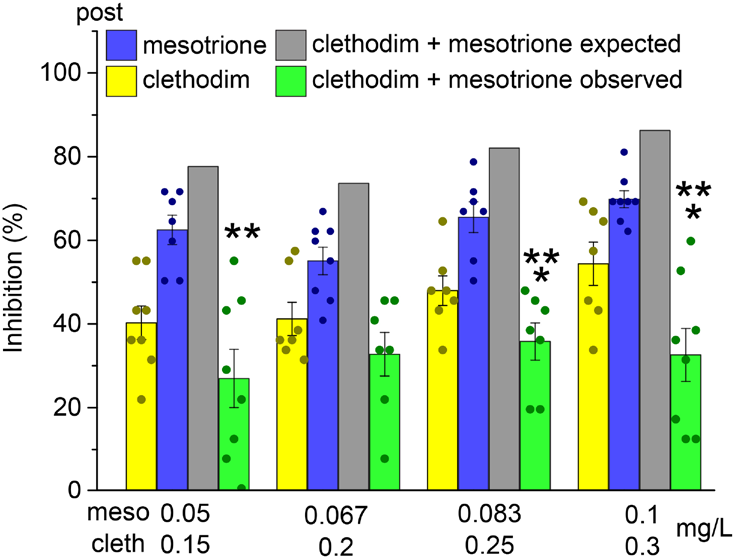
Mesotrione and clethodim were strongly antagonistic in tef. The antagonism varied between 2-fold to 3-fold when clethodim-mesotrione applied post-emergence. Error bars represent SE. Asterisks denote the significance of differences from expected for additivity * p < 0.05, ** p < 0.01, *** p < 0.001.

**Supplementary Fig. 3.**
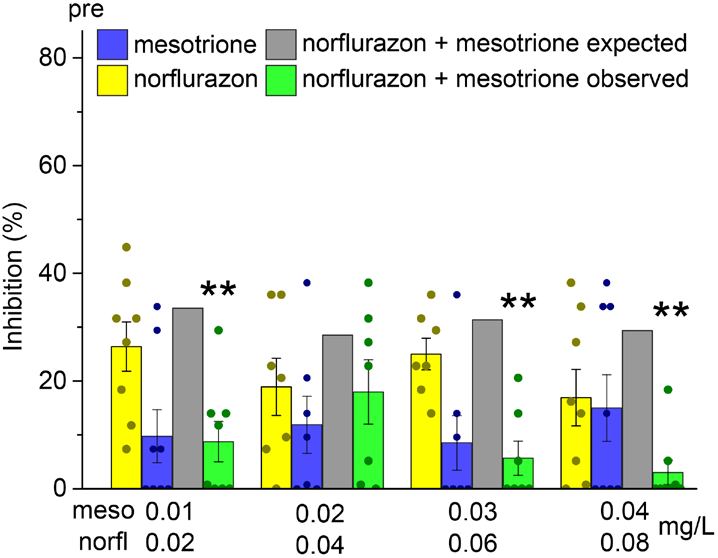
Mesotrione and norflurazon were more antagonistic in tef at higher doses. The biggest antagonism (~8-fold) was observed for mixture of 0.08 mg/L of norflurazon and 0.04 mg/L of mesotrione when applied pre-emergence. Error bars represent SE. Asterisks denote the significance difference from expected response for additivity * p < 0.05, ** p < 0.01, *** p < 0.001.

## Notes

### Competing Interest Statement

The authors have declared no competing interest.

### Summary of Updates

Following suggestions from the first pre-print on bioRxiv, we have added a figure on dose responses, shown the raw data points in all the bar charts and added a panel to one figure showing experimental design.

